# Ion-bridges and lipids drive aggregation of same-charge nanoparticles on lipid membranes

**DOI:** 10.1101/2021.11.22.468803

**Authors:** Enrico Lavagna, Davide Bochicchio, Anna L. De Marco, Zekiye P. Güven, Francesco Stellacci, Giulia Rossi

## Abstract

The control of the aggregation of biomedical nanoparticles (NP) in physiological conditions is crucial as clustering may change completely the way they interact with the biological environment. Here we show that Au nanoparticles, functionalized by an anionic, amphiphilic shell, spontaneously aggregate in fluid zwitterionic lipid bilayers. We use Molecular Dynamics and enhanced sampling techniques to disentangle the short-range and long-range driving forces of aggregation. At short inter-particle distances, ion-mediated, charge-charge interactions (ion bridging) stabilize the formation of large NP aggregates, as confirmed by cryo-electron microscopy. Lipid depletion and membrane curvature are the main membrane deformations driving long-range NP-NP attraction. Ion bridging, lipid depletion, and membrane curvature stem from the configurational flexibility of the nanoparticle shell. Our simulations show, more in general, that the aggregation of same-charge membrane inclusions can be expected as a result of intrinsically nanoscale effects taking place at the NP-NP and NP-bilayer soft interfaces.

## INTRODUCTION

The control of the aggregation of biomedical nanoparticles (NP) in physiological conditions is crucial as clustering may change completely the way they interact with the biological environment. The surface of biomedical NPs is usually designed to make NPs colloidally stable and reduce protein adsorption, to be administered via intravenous routes^1^. Once in contact with cell membranes, though, their aggregation behavior may change.

One possible way to understand and predict the aggregation behavior of NPs on lipid membranes is to look for similarities between NPs and other membrane-associated biological macromolecules. The aggregation of membrane inclusions, such as integral membrane proteins^2–4^, scaffolding proteins^5^, or toxins^6–8^, is a physical process that regulates fundamental biological functions, like endocytosis and signaling. Due to the biological relevance of aggregation processes, large efforts have been devoted to the understanding of their physical and chemical driving forces. As clearly pointed out by Johannes *et al*^9^, different attractive forces act on different length scales. Aggregation may be favored by long-range (mesoscale, 10^−9^ ÷ 10^−7^ m) membrane deformations or by direct, short-range interactions. Mesoscale effects, such as membrane curvature, capillary forces, membrane density and thickness fluctuations, are chemically non-specific. They rather depend on the size and shape of the inclusion and on its physical interaction with the bilayer^10–12^. On the contrary, short-range interactions (screened electrostatics, hydrogen bonding, van Der Waals) do depend on the chemistry of the inclusion-inclusion and inclusion-membrane interfaces.

Long range driving forces act while membrane inclusions diffuse on or within the membrane, they bring inclusions close to each other and favor the sampling of the proximity configuration space. Only at this point, short range interactions come into play to stabilize (or de-stabilize) aggregation.

There are many examples of proteins that are thought to interact via membrane-mediated forces acting within the mesoscale range, including VP1 capsid proteins^8^, Shiga toxins^6,13^, rhodopsin^3^, BAR domain proteins^14^. A common characteristic of these protein systems is rigidity. This structural feature is important, for instance, to act as a scaffold, by leaving a curvature imprint on the membrane surface; or to suppress the spontaneous membrane fluctuations, as Shiga Toxin does; or to impose membrane thickness variations through significant hydrophobic mismatch with the lipid tail region. It is because of this rigidity that continuum elastic models are successful at the quantitative prediction of aggregation regimes, taking into account the membrane elastic properties, the inclusion shape, size, and strength of membrane-inclusion coupling^15–21^. Also particle-based models with coarse grained resolution, in which the membrane is fluid but the inclusion has a rigid structure, have been successfully used to draw phase diagrams of aggregation as a function of the same physical parameters^2,4,10,11^.

At variance with proteins, though, many synthetic nanoparticles designed to interact with cell membranes for diagnostics or therapeutic purposes do *not* have a rigid interface. Most often, they couple a rigid nanoparticle core (it may be a metal, a metal oxide, a carbon-based core) to a soft shell of organic matter (covalently bound ligands, tethered polymers, physisorbed surfactants). The solid, rigid core contributes to determine the nanoparticle aspect ratio and size, which is often comparable to the size of single proteins or protein oligomers. But these characteristics can be further tuned by the soft shell, whose density, spatial extension, conformation and overall shape can dramatically change in response to the environment^22^, membranes included.

Another striking difference between biomedical nanoparticles and proteins is their surface charge. Biomedical NPs can bring large surface charges, which are useful to assure colloidal stability. The effect of surface charges on aggregation, though, is not easy to predict. On the long range, same-charge electrostatic repulsion and opposite-charge attraction are expected and modulated by the ionic-strength of the solution. On the short range, where the continuum assumptions of the Derjaguin-Landau-Vewey-Overbeek (DLVO) theory^23^ break down, same-charge attraction is also possible as a consequence of a combination of anisotropic hydrophobic interactions, charge-dipole, dipole-dipole interactions and ion bridging^24–27^. These effects can be favored by the anisotropy of NP-NP interactions, which in turn can depend both on the NP core shape and on the dynamic responsiveness of their organic shell. In Petretto *et al*.^28^, the authors show how ion bridging can be an effective short range stabilizer for the aggregation, in water, of same-charge, monolayer-protected Au NPs with an overall diameter of 5 nm. Once more, it is ligand flexibility that allows for the formation of a large number of ion bridges between the highly curved surfaces of these small NPs.

The experimental literature offers many examples^29–36^ of spontaneous aggregation of core-shell nanoparticles in contact with model lipid membranes. For these systems, though, the interplay between long range, membrane mediated forces and short range, interface mediated forces is way more difficult to disentangle than it is for rigid inclusions, both from the experimental and theoretical standpoint^22^.

Here we show, by molecular dynamics simulations, that amphiphilic, negatively charged monolayer-protected Au NPs can form large ordered aggregates upon interaction with model zwitterionic lipid bilayers. Using Molecular Dynamics (MD) at different resolutions, we show that these NPs aggregate as a consequence of both long range, membrane mediated interactions and short range electrostatic forces. We identify these contributions and provide details on their relative strength. Interestingly, different membrane mediated forces, such as curvature and lipid depletion, come into play depending on the degree of embedding of the nanoparticle in the membrane. Short-range interactions, instead, are dominated by a strong ion bridging effect, which is relevant in all the metastable states that characterize the NP-membrane interaction process. This picture emerging from simulations is consistent with the inter-particle distance measured by cryo-electron microscopy (Cryo-EM).

## RESULTS

Our reference NP is a Au NP with a core size of 4 nm, functionalized by a thiol mixture composed of the hydrophobic 1-octanethiol (OT) and the negatively charged 11-mercapto-1-undecanesulfonate (MUS), in the MUS:OT 2:1 ratio. The *in silico* models of MUS:OT 2:1 NPs, at atomistic and coarse-grained resolutions, have been presented and validated in our previous works^36–39^, and briefly described in the Methods section. Here, we recall a peculiar feature of the interaction between these NPs and fluid zwitterionic lipid membranes. The NP-bilayer interaction evolves through 3 different metastable states, corresponding to different degrees of embedding of the NP into the membrane: adsorbed, semi-snorkeled and fully snorkeled, as shown in Figure S1. These metastable states are long-lived. Both adsorbed and snorkeled NPs can be observed experimentally, as reported in^30,31^. In the following sections, we address the aggregation behavior of MUS:OT 2:1 Au NPs in water and upon contact with a DOPC bilayer. When in contact with DOPC, we will describe the different aggregation mechanisms that take place depending on the degree of NP embedding in the membrane.

### Ion bridging drives the aggregation of NPs in water

As a rule of thumb, aggregation of MUS:OT NPs should be inhibited by the repulsion of the negatively charged sulphonate terminals; yet, as shown in Petretto *et al*.^28^, there is computational and experimental evidence that dimers and other small aggregates of NPs can be observed at low ion concentrations. Cryo-EM images and simulation^28^ for MUC:OT and MUS:OT NPs with core diameters in the 2-5 nm range show that the NP-NP distance is compatible with a configuration in which the MUS ligands are extended (see Figure 1a).

**Figure 1:**
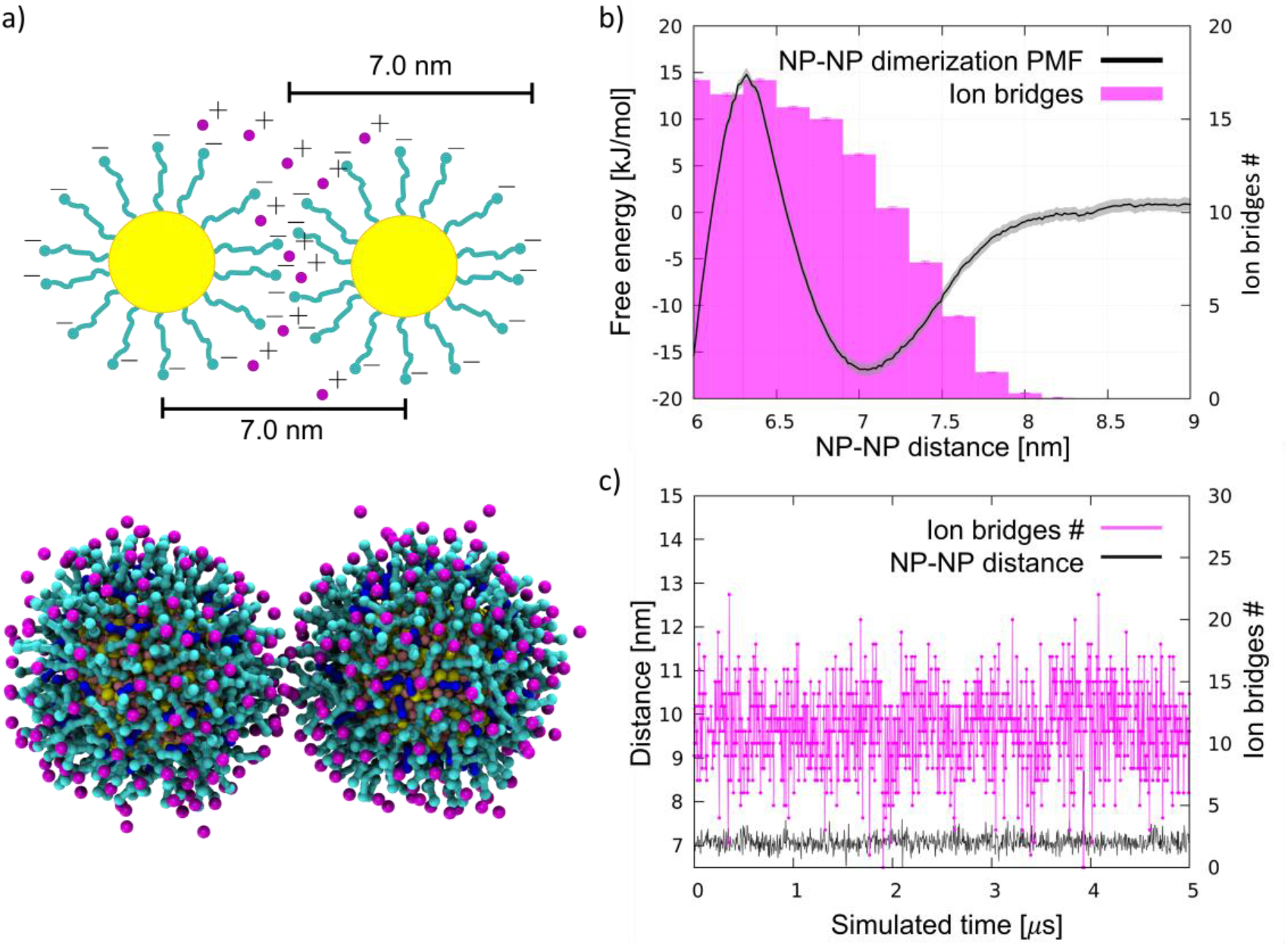
NP aggregation in water. a) Top: a sketch of an NP-NP dimer in water, in which the MUS ligands (cyan) are extended, and their negatively charged terminals, at the NP-NP interface, are bound by ion bridges (pink). Bottom: the same configuration imaged during an unbiased coarse-grained MD run. Au core is yellow, MUS ligands are cyan, OT ligands are blue, and counterions are pink. Water is not shown for clarity. b) Dimerization free energy profile (black) and corresponding average number of ion bridges (pink histogram) as a function of the distance between the NP centers of mass. The dimerization profile is cut at 6 nm. The complete profile, showing the energy well corresponding to hydrophobic NP-NP interaction, in shown in Figure S2c. c) Number of ion bridges as a function of time for the unbiased simulation containing 2 NPs in water.

Here, we tested the ability of our coarse-grained Au NP model to capture the ion bridging effect. We simulated by unbiased MD the spontaneous aggregation of MUS:OT 2:1 NPs with a core diameter of 4 nm. We performed 3 unbiased simulations with 2, 10, and 20 NPs in water and NaCl 150 mM, plus counterions. In all runs, within short simulation times, the NPs aggregate forming a cluster; clusters appear to have a rather dynamic shape but are stable throughout the 10 µs simulations. Moreover, the NP-NP distance is peaked at 7.0 nm, as shown by the radial distribution functions (Figure S2a). This distance corresponds to NPs interacting in an extended-ligand conformation.

Since the NPs interact through the negatively charged sulphonate ligand terminals, the most plausible explanation for the stabilization of the NP-NP contact is the presence of ions that bridge the charged beads of the two different NPs. The analyses of the MD runs confirm this hypothesis. For the 2 NP simulation, Figure 1c shows the number of ions that are in contact with both NP simultaneously; on average, about 10 ion bridges can be identified. The number of ion bridges undergoes significant fluctuations, pointing to a quite dynamic NP-NP interface.

To characterize the thermodynamic features of this interaction, we calculated the potential of mean force (PMF) of dimerization using Umbrella Sampling^40^ (US), as described in the Methods section. The complete profile, shown in Figure S2c, is characterized by two minima and is qualitatively similar to the dimerization PMF of the MUC:OT NPs described in Petretto *et al*^28^. The deepest minimum, lying at an NP-NP distance of 5.2 nm, corresponds to the hydrophobic contact, in which the hydrophobic stretches of the NP ligands face each other, as better detailed in Petretto *et al*^28^. and shown in Figure S2c. The second minimum, shown in Figure 1b and located at a NP-NP distance of about 7.0 nm, corresponds to the same dimer configuration we sampled in the unbiased simulations. This energy minimum, which is about 1 nm wide, reflects the short range nature of ion bridging, as confirmed by the plot of the number of ion bridges as a function of the NP-NP distance (Figure 1b). A significant free energy barrier prevents the dimer from collapsing into the hydrophobic contact stage within the duration of our unbiased CG runs.

We further verified the reliability of our coarse grained approach by means of simulations performed with the united atom OPLS-ua^41^ force field (details in the Methods section). In OPLS-ua unbiased simulations, we recorded an equilibrium NP-NP distance of 6.8 nm, combined with a number of ion bridges in excellent agreement with those observed in the coarse grained simulations, as reported in Figure S3.

We remark that, *a priori*, our CG approach is expected to provide only qualitative indications about NP aggregation in water. Indeed, at CG level electrostatic interactions are cut-off at rather short distances (1.1 nm). Moreover, the description of ions is a critical aspect of both coarse grained and atomistic force fields^42,43^. The reader can find more details on the comparison between the OPLS-ua simulations and the CG ones, and a critical discussion on the intrinsic limitations of the coarse-grained approach, in the ESI. *A posteriori* and despite these caveats, we find a satisfying agreement between the coarse grained and the united-atom simulations, as well as between the CG simulations and the experimental data that will be presented in the next section.

In the following sections, the simulations of the aggregation of NPs in contact with lipid bilayers will be performed at the coarse-grained level, which is the only approach that allows for sampling the relevant time and length scales of the aggregation process in the bilayer environment. Validations at united atom resolution will be explicitly referred to whenever performed.

### Ion bridging drives the aggregation of NPs adsorbed on membranes

Cryo-EM images show large aggregates of MUS:OT NPs on the surface of DOPC liposomes, as shown in refs^30,31,44^ and in Figure 2. The aggregates are located both at the interface between adjacent liposomes^31,44^ and on their free surfaces, and can be formed by NPs that are simply adsorbed on the bilayer surface, or partially or fully embedded in it^31,44^. In the large aggregate of MUS:OT NPs shown in Figure 2b, the average distance between neighbor NPs was calculated as 6.12 ± 0.5 nm. This distance is much larger than the NP diameter, suggesting that sulfonated ligands of the interacting NPs are in an extended configuration, parallel to the membrane plane.

**Figure 2:**
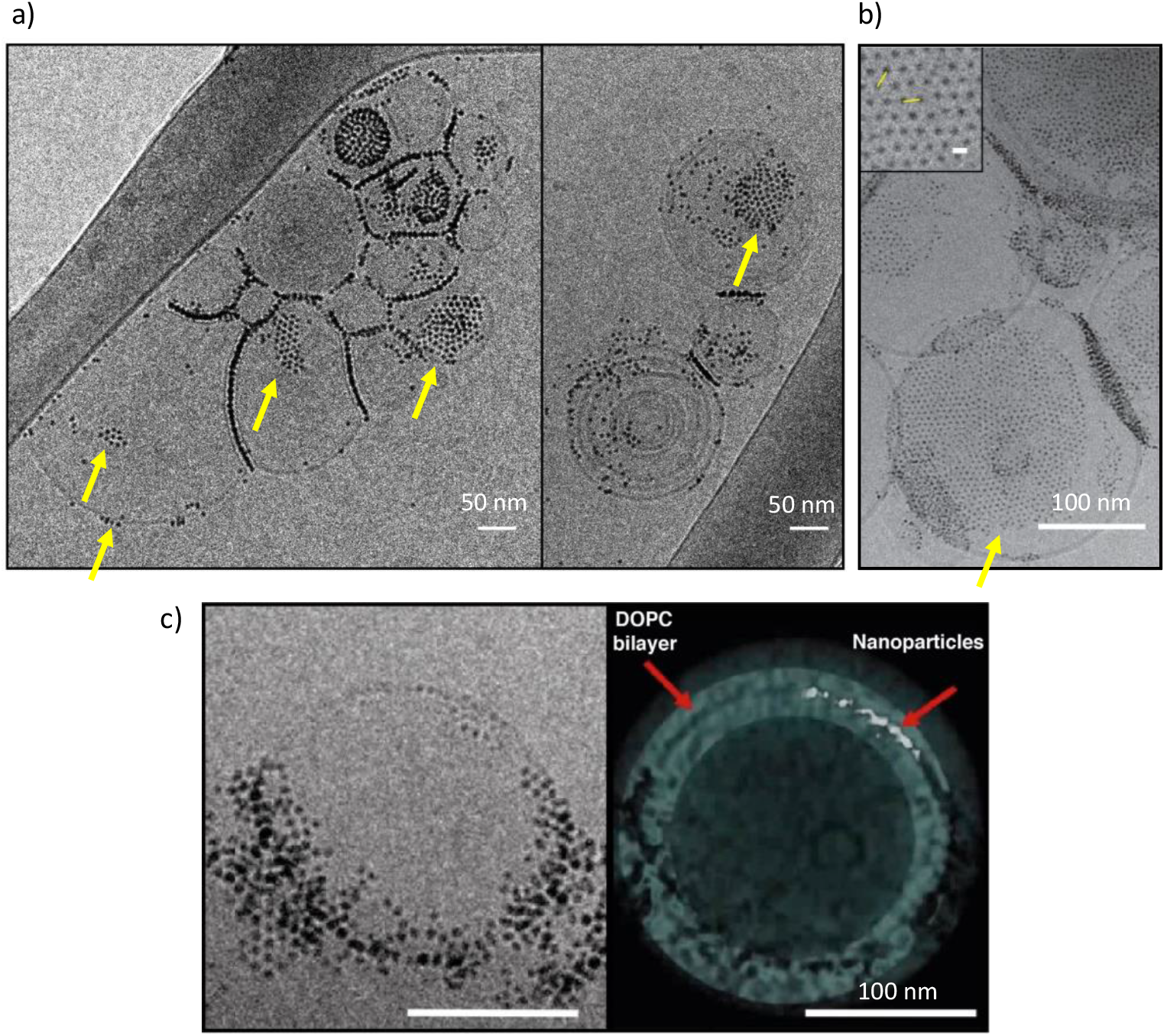
Cryo-EM imaging of large, ordered aggregates of MUS:OT NPs in contact with DOPC liposomes. a) aggregates of MUS:OT NPs (30% OT content) with a core diameter 5 ± 0.9 nm, on large DOPC liposomes; b) aggregates of MUS:OT (10% OT content) NPs, with a core diameter of 1.7 ± 0.5 nm, on large DOPC liposomes. The average distance between neighbor NPs was calculated as 6.12 ± 0.5 nm, suggesting that sulfonated ligands of the interacting NPs are in an extended configuration, parallel to the membrane plane. c) A frame from aligned file of tomogram acquisition and its 3D reconstruction. The nanoparticles, MUS:OT 30%OT (5 ± 0.8 nm) are aggregated and the 3D reconstruction suggests they are embedded within the membrane core. A description of the NP synthesis and of the imaging set up can be found in the ESI.

We start our computational investigation with the analysis of NPs that are simply adsorbed, not embedded, on the DOPC bilayer. This is the first metastable state of the interaction between the NP and the membrane. In this configuration, the sulphonate ligand terminals get in contact with the zwitterionic heads of DOPC lipids (Figure S1a). Two different pathways, possibly coexisting, can lead to the formation of such large adsorbed aggregates. The first is the diffusion of isolated, adsorbed NPs on the bilayer surface. The second is the flattening of tridimensional NP aggregates, pre-formed in water, upon contact with the bilayer.

#### Aggregation upon diffusion of single adsorbed NPs

The effect on membrane curvature of single adsorbed NPs is relatively small^44^ (Fig S4a), and it is reasonable to expect that, upon diffusion, adsorbed NPs may aggregate via a similar aggregation mechanism as that observed in water, the only difference being the planar constraint.

We simulated systems with 2 and 9 NPs adsorbed on the membrane. The snapshot of Figure 3a shows the dimer spontaneously formed during the simulation. The dimer was stable for most of the simulated time and, once again, stabilized by ion bridges, as shown in Figure 3b. Further confirmation that dimerization is due to ion bridging is provided by the PMF of Figure 3c, which is very similar to the one obtained in water. In the 9 NP simulation, we initialized the run by placing the 9 NPs far from each other. The NPs spontaneously aggregated within 20 µs, forming a bidimensional aggregate with a roughly hexagonal packing (Figure 3d). The NP-NP first neighbor distance within the larger aggregate is again around 7.0 nm (Figure S4b), and it corresponds to an extended conformation of the ligands in the shell of the NPs. These data match very well the close-packed arrangements observed experimentally, as shown in Figure 2b and in refs.^30,31^.

**Figure 3:**
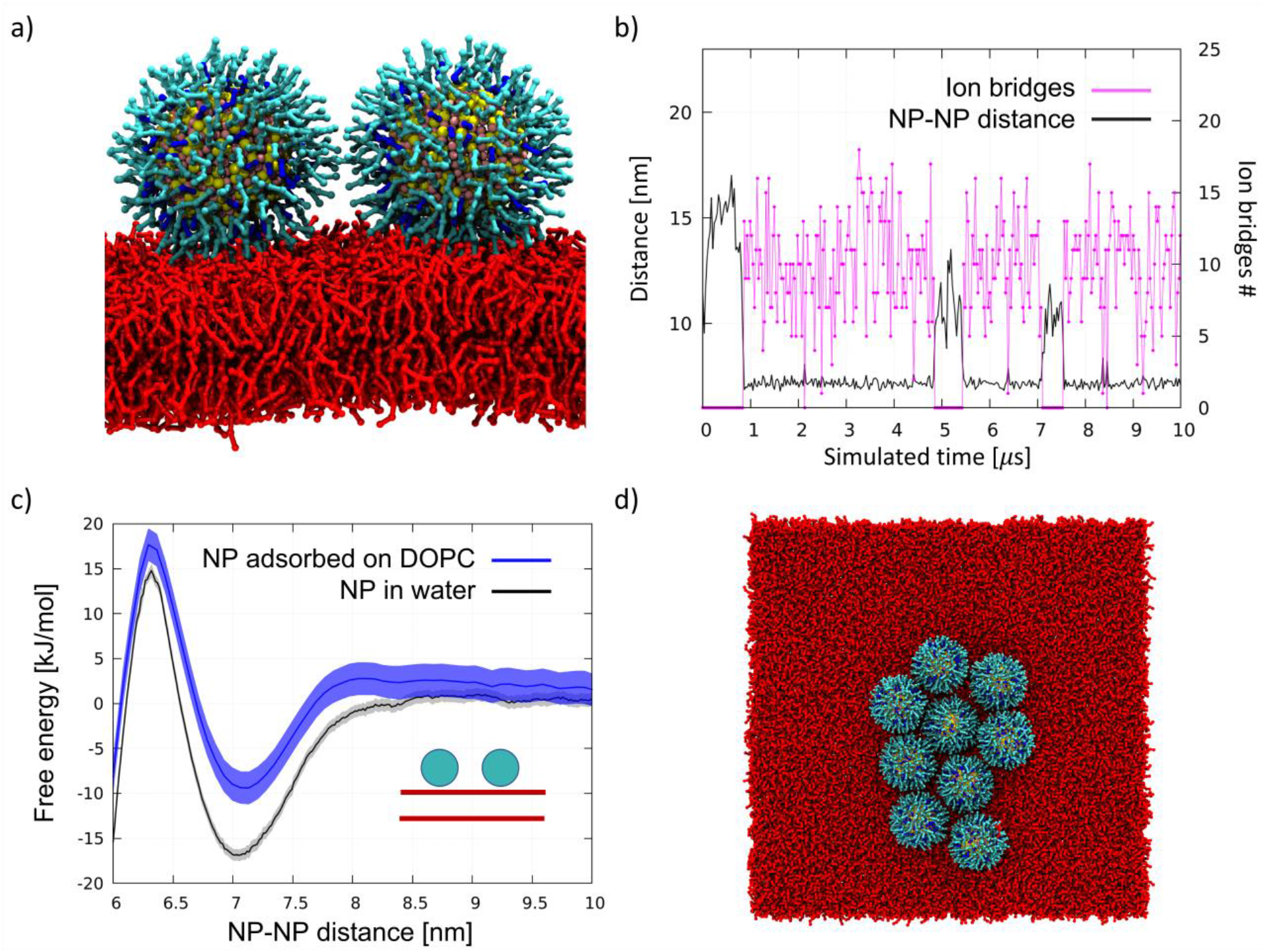
Aggregation of DOPC-adsorbed NPs. a) Lateral view of the NP dimer adsorbed on DOPC. Same color code as in Figure 1, DOPC in red. Water and ions are not shown; b) Number of ion bridges (pink) and NP-NP distance (black) in the simulation of 2 adsorbed NPs diffusing on the DOPC membrane surface; c) Dimerization PMF; the dimerization profile on DOPC is similar to that obtained n water; d) Top view of the aggregate spontaneously formed upon diffusion of 9 NPs adsorbed on the DOPC membrane surface.

#### Flattening of a tridimensional aggregate, previously formed in water

We simulated a system with a tri-dimensional aggregate of 6 NPs, placed in water, and a planar membrane patch. Within the simulated 4 µs, we observed the cluster flattening on the membrane (Figure S4c).

Our data suggest that both pathways, namely the diffusion of isolated, adsorbed NPs and the flattening of 3D NP aggregates upon interaction with the bilayer, can lead to the formation of large bidimensional aggregates of adsorbed NPs, which are then stabilized by ion bridging.

### A complex interplay of electrostatics and membrane deformations drive the aggregation of membrane-embedded NPs

Cryo-EM images show single NPs embedded in the membrane core and even the presence of large NP aggregates constituted by membrane-embedded NPs, as shown in refs.^30,44^ and recalled in Figure 2c. In the following, we address the simulation of the aggregation of NPs in the fully- and semi-snorkeled configurations.

#### Fully snorkeled NPs: ion bridging and lipid depletion drive NP aggregation

In this section, we describe aggregation upon diffusion of single, fully snorkeled NPs. We ran 3 unbiased simulations with 2, 9, and 36 NPs, observing, in all cases, spontaneous aggregation, as shown in Figure S5a. In figure 4a we show the PMF of dimerization of fully snorkeled NPs, compared with the PMF of NPs in water. The minimum of the former is shifted to slightly larger distances, about 7.8 nm; again, we observe the formation of ion bridges (details in the Supporting Information, Figure S5b).

**Figure 4:**
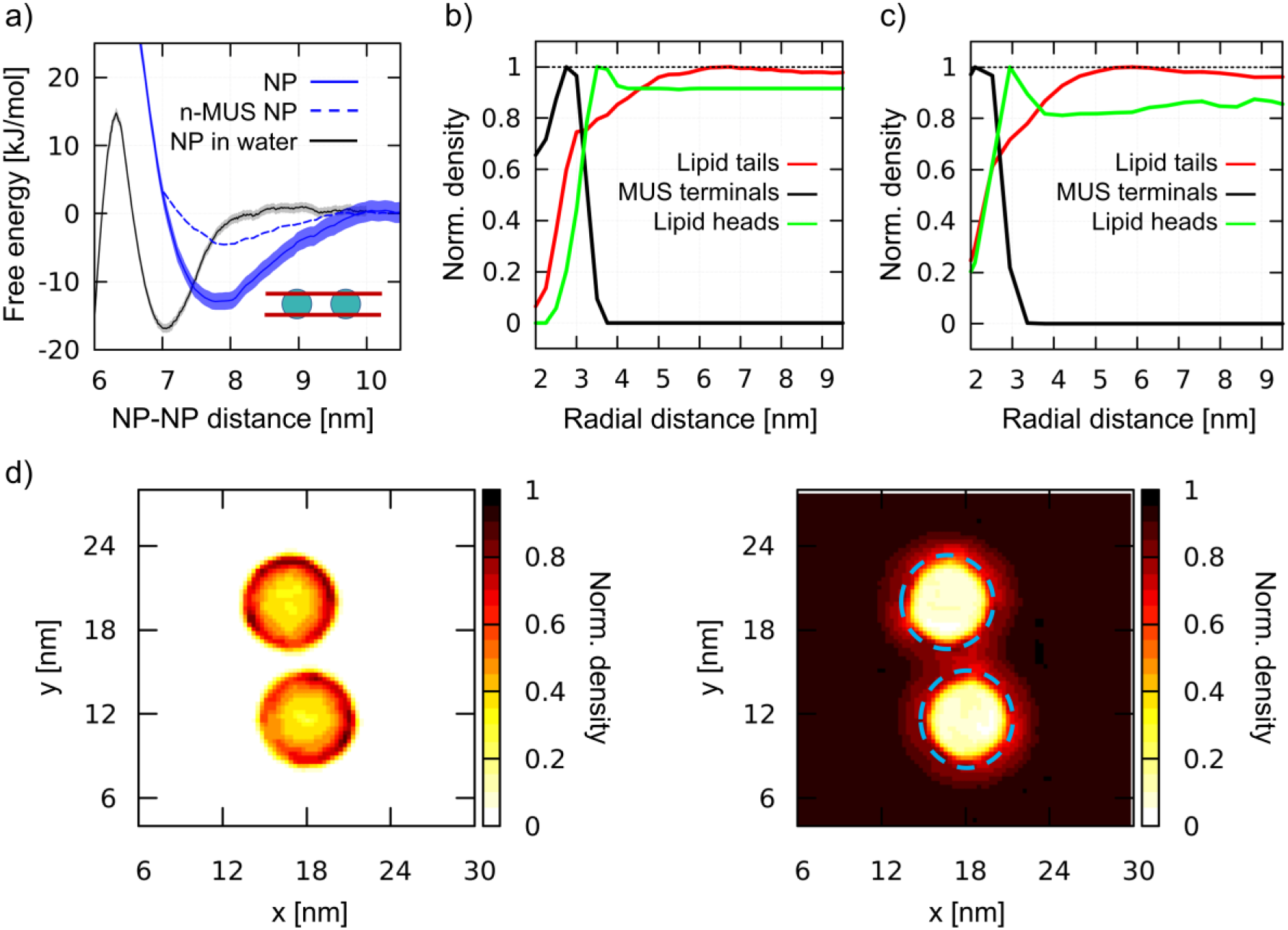
Aggregation of fully snorkeled NPs. a) Dimerization PMF. Fully snorkeled NPs in DOPC (blue, solid line); fully snorkeled NPs with n-MUS ligand terminals in DOPC (blue dashed line, error bars are not shown but they are comparable to those of regular NPs); NPs in water (grey, from Figure 1). b),c) Normalized density of charged MUS ligand terminals (black), lipid tails (red), and lipid heads (green) as a function of the distance from a single, fully snorkeled NP in DOPC, calculated at coarse-grained (b) and united atoms (c) resolutions. d) Normalized number density maps of MUS ligands (left) and DOPC tails (right) around a fully snorkeled NPs dimer; the blue dashed circle on the right corresponds to the external edge of the MUS corona, as measured from the left map.

However, quite interestingly, the dimerization profile of fully-snorkeled NPs shows that the interaction is now a long range one. The PMF starts decreasing at about 10 nm distance: at this distance, the ion bridging effect cannot be present; hence the membrane must have a role in promoting dimerization.

To isolate and identify the membrane contribution to aggregation, ruling out ion bridging and other electrostatic effects, we prepared a modified version of the Martini NP topology in which the charge of the sulphonate ligand terminal is neutralized. We will refer to these NPs as n-MUS NPs. The unbiased simulations of n-MUS NPs show that the aggregation is still present, albeit less stable. Indeed, as shown in Figure 4a, the corresponding dimerization profile is similar to the one of the regular NPs, with the same long range attraction but a shallower minimum (∼5 kJ/mol against 15 kJ/mol). This simple test confirms that the long-range driving force for aggregation does not depend on electrostatics but is instead a membrane-mediated interaction effect.

To determine how the membrane induces this long-range attraction, we analyzed the membrane structure around a single, fully snorkeled NP. The fully snorkeled NP does not alter membrane curvature (Figure S6a); membrane thickness is slightly reduced nearby the NP, but this is a short-range effect (< 6 nm from the NP center). Hence, we can rule out curvature effects and hydrophobic mismatch from the possible causes of the observed long range interaction.

Interestingly, the analysis of the density of the lipid tails around the NP showed an area of reduced density with a spatial extension that overlaps well with the attraction basin of the PMF. Figure 4b shows the radial distribution function of MUS ligand terminals, DOPC heads, and DOPC tails with respect to the NP center. The lipid tails’ density is depleted in a circular region with a radius of about 5 nm, extending more than 2 nm beyond the MUS terminals. This result is also confirmed at united atom resolution (Figure 4c and Figure S6d), where we observe the same lipid tail depletion range. When 2 NPs approach each other, their lipid depleted auras start overlapping at a NP-NP distance of 10 nm. This distance is precisely the onset of dimerization, as shown by the PMF profiles (Figures 3a). 2D maps of the normalized number density of MUS ligand terminals and DOPC tails around a dimer, reported in Figure 4d, show the overlap of the lipid depletion auras when the NPs are closer than 10 nm. The lipid depletion aura is also observed in the neutralized system (n-MUS:OT NPs, Figure S6b). In this case, the range of depletion is similar, but its magnitude is lower, coherently with the shallower PMF minimum (Figure 4a, dotted line).

#### Semi-snorkeled NPs: ion bridging and membrane curvature drive NP aggregation

For this intermediate state, the dimerization process and the subsequent formation of aggregates is more complex. We ran unbiased simulations of 2 and 9 semi-snorkeled NPs, initially far apart from each other. In the 2 NP simulation we observe again the spontaneous formation of a dimer. Figure 5a shows the PMF of dimerization of semi-snorkeled NPs and the number of ion bridges that form while the 2 NPs are approaching each other. The dimer minimum (at about 7.5 nm) is thermodynamically less stable than in the adsorbed and fully snorkeled case. Moreover, the dissolved state is separated by the dimer state by a small free energy barrier (about 5 kJ/mol) that was absent for fully snorkeled NPs. In the 9 NP unbiased simulation, the aggregation is slower than for fully snorkeled NPs, due to the presence of the aforementioned free energy barrier. Nevertheless, we observe again the formation of a stable aggregate, with features similar to the aggregate formed in the fully snorkeled and adsorbed cases: ion bridges mediate NP-NP interactions within the aggregate.

**Figure 5:**
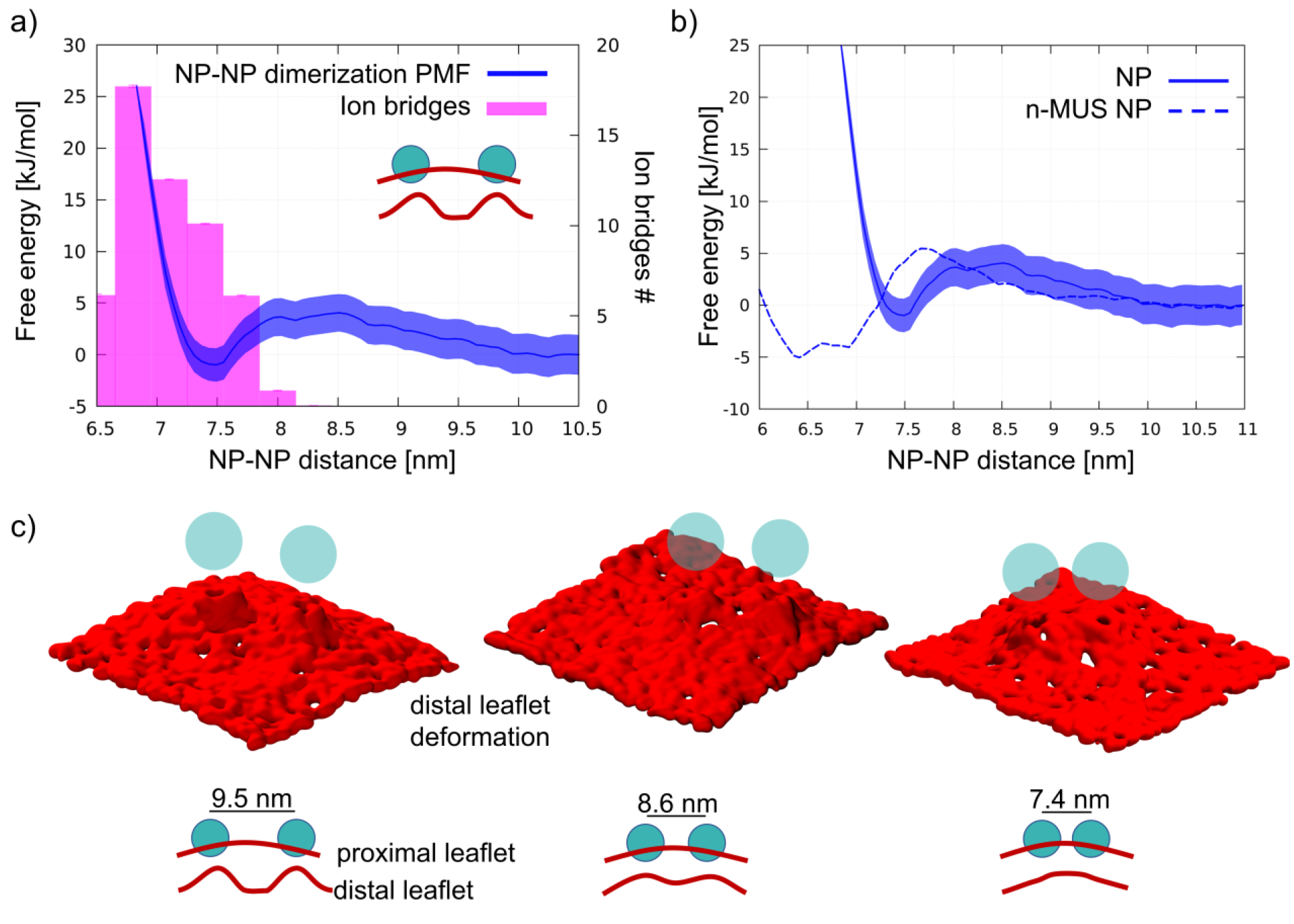
Aggregation of semi-snorkeled NPs. a) Dimerization PMF (solid blue line) and average number of ion bridges (pink histogram). b) Dimerization PMF as in a), compared to the dimerization PMF of n-MUS NPs (dashed blue line, error bars are not shown but they are comparable to those of regular NPs); c) Surface representation of the lipid headgroups of the distal leaflet (red surface). When the semi-snorkeled NPs do not interact with each other (9.5 nm), each NP heavily deforms the distal leaflet. As the NPs approach each other, their deformations start overlapping (8.6 nm) until the deformations collapse into a single, smoother deformation (7.4 nm, dimer configuration).

While ion bridging certainly contributes to stabilize aggregation, the membrane deformations induced by semi-snorkeled NPs are more pronounced than those caused by fully snorkeled NPs. Indeed, the membrane is significantly curved by both single and aggregated semi-snorkeled NPs, as shown in Lavagna et al^44^ and in Figure S1b. Therefore, we compared the behavior of regular and neutralized n-MUS NPs in the semi-snorkeled configuration to verify whether and how membrane curvature effects can contribute to the aggregation process.

We ran an unbiased simulation in which 9 n-MUS NPs, in the semi-snorkeled configuration, were free to diffuse in a planar membrane patch. The NPs aggregate again, but the NP clusters are less stable than in presence of charged ligands, and rearrange dynamically over the time scale of the simulation. Figure 5b shows a comparison between the PMF of dimerization of charged and neutral semi-snorkeled NPs. The NP-NP distance corresponding to the lowest free energy decreases from 7.5 to 6.6 nm. The n-MUS NPs PMF shows the same free energy barrier (about 5 kJ/mol) observed for the regular NPs. This barrier is responsible for the slower aggregation kinetics. As electrostatics can not be responsible for this repulsion, this effect must be accounted for by membrane deformations. The analysis of the lipid tail density around the semi-snorkeled NPs shows that the NPs do not induce significant density perturbation. However, the curvature of the membrane during the dimerization process appears to be significantly perturbed, especially in the distal leaflet. In Figure 5c we show 3 snapshots representative of the conformations of the distal leaflet lipid heads at the NP-NP separation of 9.5, 8.6, and 7.4 nm. In the 9.5 nm snapshot, the two distinct NP-induced deformations are well distinguishable and separated, and the membrane is flat in between; as the two NPs get closer to each other, the two deformations start overlapping, and in the final snapshot, at 7.4 nm, a single deformation, characterized by a smoother curvature profile, is formed. The same behavior can be observed for the charged NPs. The collapse of the two deformations into a single, smoother deformation is the process that requires overcoming the free energy barrier observed in the PMF.

We can conclude that the dimerization profile of semi-snorkeled NPs derives from the superposition of two effects, acting at slightly different distances. The repulsive barrier due to the approaching membrane deformations contrasts dimerization, while the formation of ion bridges favors it. As a result, the dimerization range of semi-snorkeled NPs is narrower and less thermodynamically favorable than for adsorbed and fully snorkeled NPs.

## CONCLUSIONS

In this paper, we have characterized the aggregation of monolayer-protected, anionic and amphiphilic Au NPs, with a core diameter of 4 nm, in contact with DOPC membranes. We found that aggregation results from three main contributions. The first is ion-bridging. Ions are the leading driving force to aggregation of adsorbed NPs. In this configuration, the NP-NP interface is similar to that of solvated NPs, and the result is in line with those obtained by Petretto *et al*^28^. When the NPs interact with the membrane via hydrophobic contacts, either in the fully- or semi-snorkeled configuration, ion-bridging is still present at short-range, but two other aggregation forces come into play, which are membrane-mediated. For NPs in the fully-snorkeled configuration, lipid depletion around the NP gives a significant contribution to aggregation, an effect that has a remarkably long range. For NPs in the semi-snorkeled configuration, it is the minimization of the NP-induced membrane curvature to drive aggregation.

Due to the slow kinetics of interaction between these amphiphilic Au NPs and phosphatidylcholine membranes, all the intermediate, metastable states of NP-membrane interactions are relevant. In a recent paper, we showed how amphiphilic Au NPs can induce different curvatures on the bilayer, depending on their degree of embedding^45^. When aiming at an interpretation of the effects of ligand-protected NPs on membranes, it is crucial to take into account how different configurations of their ligand shell may change the structure of the NP-membrane complex, and how dynamic the shift from one configuration to the other can be.

In this paper we gained a deep, one of a kind understanding of aggregation for MUS:OT NPs in DOPC membranes. Our results, though, are not system-specific, and our claims have a quite general validity. First, aggregation between same-charge nanoscale membrane inclusions can be expected, even when their surface charge density is large. This result may hold for a broad range of engineered NPs, whose fate in the biological environment should be controlled; and it could even hold for accidental NPs, whose fate in the biological environment should be understood and predicted. Second, the NP ligand flexibility is a key descriptor of the NP physico-chemical nature, and it should be carefully taken into account when interpreting experimental data related to NP-membrane interactions, as already pointed out for NP-protein interactions^46^. Indeed, when the membrane inclusion has an amphiphilic, flexible surface, such an interface adapts itself to its environment as the interaction with the membrane unrolls. This means that very different mechanisms of aggregation may be activated at the same time or in close sequence, due to the dynamic configurational changes of the inclusion-membrane interface. Last but not least, we envisage that these results will seed further exploration of the possible differences and analogies between ligand protected NPs and proteins. Thanks to the increasing ability to engineer NPs with the desired surface properties, and to the fundamental understanding of their behavior in the biological environment, NPs are gaining recognition as models to interpret protein-protein and membrane-protein interactions. Our conclusions may be especially relevant in those cases when protein aggregation is attributed to specific bridging interaction with ions^47–50^, not only in solution but also in the membrane environment^51^.

## METHODS

### Models

#### Coarse grained (CG) model

All the simulations relied on the standard Martini force field^52,53^ for both membrane and ligands. The NP model was custom developed and already used in previous works^36,38,39,54^; it is modeled as a 4 nm diameter hollow sphere composed of 346 Au and 240 uniformily distributed S atoms. A total of 240 ligands (168 MUS, 72 OT) are bound, in random order, to the S atoms of the core. NP topologies of the NPs can be found at this <u>link</u>.

#### Atomistic model

As atomistic validation for some of the results obtained with the CG model, we also ran simulations using the united atom OPLS force field^41^ and the rigid SPC/E water model^55^. The core of the NP is the same as in the Martini model, while the ligands are mapped using the united atom OPLS interaction parameters. Topologies of the nanoparticles are available at this <u>link</u>.

### Simulation setup

All simulations were run with Gromacs 2020.4, using the leapfrog integrator and the NPT ensemble, at a temperature of 310 K and a pressure of 1.0 bar. For temperature coupling, we used the velocity rescale^56^ algorithm. For pressure coupling, we used the Parrinello-Rahaman^57^ algorithm in production runs, and the Berendsen^58^ algorithm in equilibration runs.

#### CG simulations

In all CG simulations, we used a time step of 20 fs, and we treated the electrostatic interactions using a cut-off of 1.1 nm^59^. We set the temperature coupling constant *τT* to 1 ps. Concerning the pressure coupling, we used a *τP* =12 for production runs and *τP* =4 for equilibration runs. We set the compressibility at 3·10^−4^ bar^-1^. For membrane simulations, the pressure coupling was semi-isotropic to ensure vanishing surface tension in the bilayers.

#### Atomistic simulations

We used a 2 fs time-step in atomistic simulations, with both coupling constants *τT* and *τP* set to 1 ps. The compressibility of the systems was set to 4.5·10^−5^ bar^-1^. Since all the atomistic simulations involved systems without membranes, we used the isotropic pressure coupling scheme.

#### Free energy calculations

In order to obtain the Potential of Mean Force (PMF) profiles, we used the umbrella sampling method^60^ with the Gromacs wham tool^61,62^ to estimate errors. We used 0.2 nm spaced windows and a force constant for pulling of 750 kJ mol^-1^ nm^-2^. Pulling rates were always zero. Bootstrap analysis was performed with 100 bootstrap samples and tolerance set at 1·10^−6^.

All bilayers were formed using the insane^63^ tool or adapted from previous simulations. A list of simulated systems, as well as the description of all the initial configurations used for the MD runs, are reported in the Supporting Information.

## ASSOCIATED CONTENT

### Supporting Information

The file Lavagna_SI_NPaggregation.docx is available free of charge.

## AUTHOR INFORMATION

### Author Contributions

GR conceived the idea and discussed it with FS, DB and EL. EL performed most of the MD simulations and free energy calculations, with contributions from DB and ALDM. EL and DB analyzed the data with the supervision of GR. ZPG performed the NP synthesis and Cryo-EM imaging under the supervision of FS. EL and DB wrote a first draft of the manuscript, which was finalized with contributions of all authors. All authors have given approval to the final version of the manuscript.

## ACKNOWLEDGMENTS

GR acknowledges funding from the H2020 ERC Starting Grant BioMNP – 677513. GR, EL and DB acknowledge funding by MIUR – DIFI Dipartimento di Eccellenza 2018-2022 for computational resources. The authors thank Stefano Vanni, Emanuele Petretto, and Quy K. Ong for many fruitful discussions, for sharing their results and for a critical reading of the manuscript.

