## Supplemental Information for "Ion-bridges and lipids drive aggregation of same-charge nanoparticles on lipid membranes"

***List of simulated systems***

*NPs in water.* For Martini systems comprising 10 or 20 NPs in water without the membrane, we placed the NPs at a fixed distance on a 3d grid, in a cubic box of 36 nm side. Dimer simulations (both biased and umbrella sampling) were run in a smaller water box of 20 nm side. All the unbiased simulations were first equilibrated for 100 ns. For the OPLS dimers, we used a rhombic dodecahedron box with an 18 nm distance between periodic images. The two NPs were placed at a 7 nm distance to reproduce the free energy minimum configuration visible in CG simulations. We ran simulations at 150 mM and 300 mM concentrations, and, for each concentration, we used $Na^+$ counterions and $Ca^{2+}$ counterions for a total of four 100 ns simulations.

*Adsorption of NP aggregates on the membrane.* For these simulations, we used a 6 NP aggregate extracted from an unbiased simulation with only water and placed it in a 38x28x26 $nm^3$ box together with a formed DOPC bilayer with 4050 lipids.

*Adsorbed NPs simulations.* The box was 47x47x12 nm3 and contained a 5616 DOPC bilayer. We placed the 9 NPs at the adsorption distance from the bilayer, all on the same side and on a 3x3 grid.

*Semi- and fully snorkeled NPs simulations.* In this case, we first prepared a system with a single NP snorkeled in the membrane and then multiplied it in a 9x9 grid to obtain the entire configuration. In order to create a membrane patch with an embedded NP, we used the insane tool[54], placing the NP at a relative distance of 0 nm (fully snorkeled) or 3 nm (semi-snorkeled) compared to the bilayer midplane. We then equilibrated the system freezing the NP core in order to let the membrane-NP system stabilize. The final box (39x39x16 $nm^3$) contained 9 NPs and 4158 DOPC lipids.

*Lipid-depletion simulations*. We conceived a particular setup to better analyze the lipid depletion auras around NPs and produce the maps of Figure 4. Indeed, these simulations have been performed in the NVT ensemble (after NPT equilibration) and keeping the Au cores of the NPs frozen in the XY plane to reduce the noise coming from the simulation box oscillations and the NP movements (migration/rotation) inside the membrane. The freezing in the XY plane was essential for the dimer simulations to keep the two NPs at a fixed distance. All the ligands were always free to move, and we verified that the same effect (lipid tails depletion), although appearing noisier, was present also in the absence of core freezing.

*n-MUS simulations.* In these cases, the starting configurations were taken from the respective system's starting configurations with standard NPs.

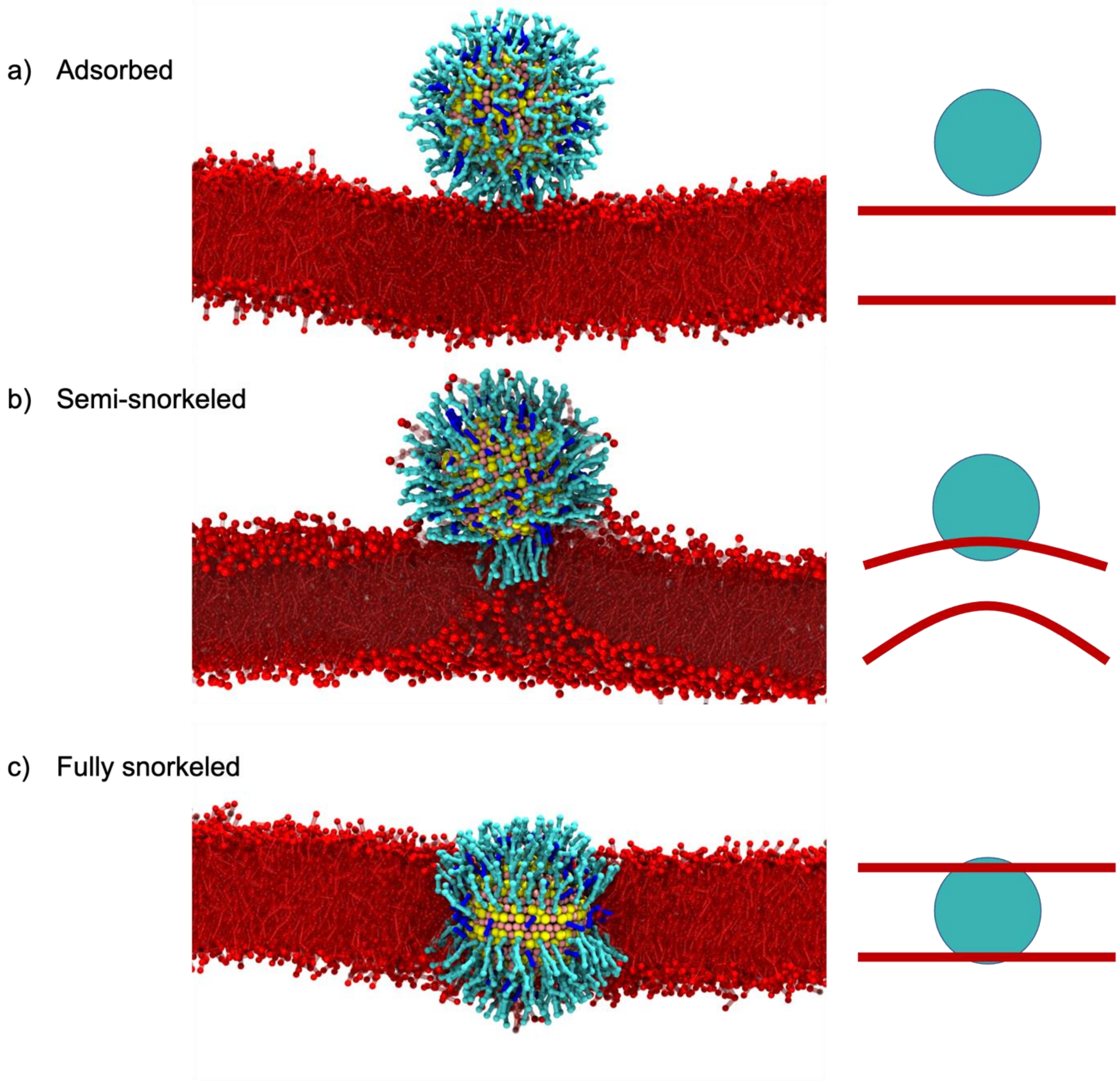


**Figure S1 -** The NP-bilayer interaction evolves through 3 different metastable states, corresponding to different degrees of embedding of the NP into the membrane: a) adsorbed, b) semi-snorkeled and c) fully snorkeled. These three metastable states are long lived and can be observed experimentally, by cryo-EM. As the NP penetrates the membrane core, it induces different curvatures to the bilayer, as discussed in Lavagna et al.[17] and shown by the sketches on the right.

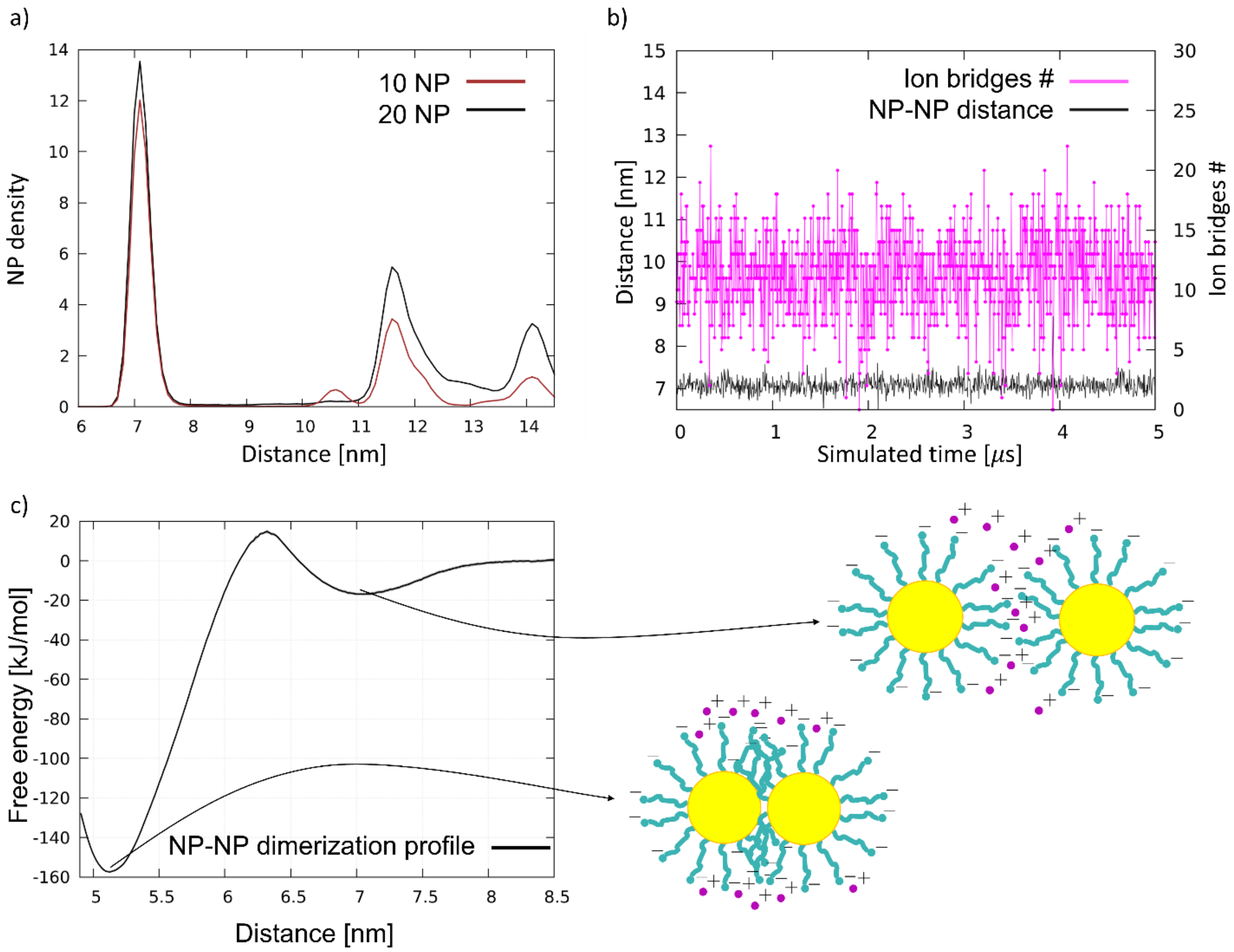


**Fig S2. Aggregation in water** a) Rdf of NP-NP distances for the unbiased simulations with 20 NP (black) and 10 NP (brown). b) Ion bridges count (magenta) and NP-NP distance (black) in the 2NP simulation. c) Dimerization PMF profile of the NP in water, with a sketch of the relative NP-NP position in the two minima: the hydrophobic contact (5.1 nm) and ion bridging (7.0 nm).

**NP aggregation in water.** In addition to the ion bridged dimerization state, thoroughly discussed in the main text, two NPs can stabilize their interaction in a state of hydrophobic contact, in which the two ligand shells interpenetrate, and the cores are almost in contact. Within the time scale of our unbiased CG simulations, NPs cannot cross the energy barrier that leads to NP-NP hydrophobic contact. The PMF (Figure S1c) shows that the ion bridging energy well is separated from the hydrophobic contact state, which is represented by the energy minimum at 5.2 nm, by a barrier of about 35 kJ/mol and from the dissolved state by a barrier of about 18 kJ/mol. These barriers are large enough to explain the stability of the dimer in this configuration throughout the simulated timescale, and also to explain the experimental observation of both the stable and metastable aggregation states.

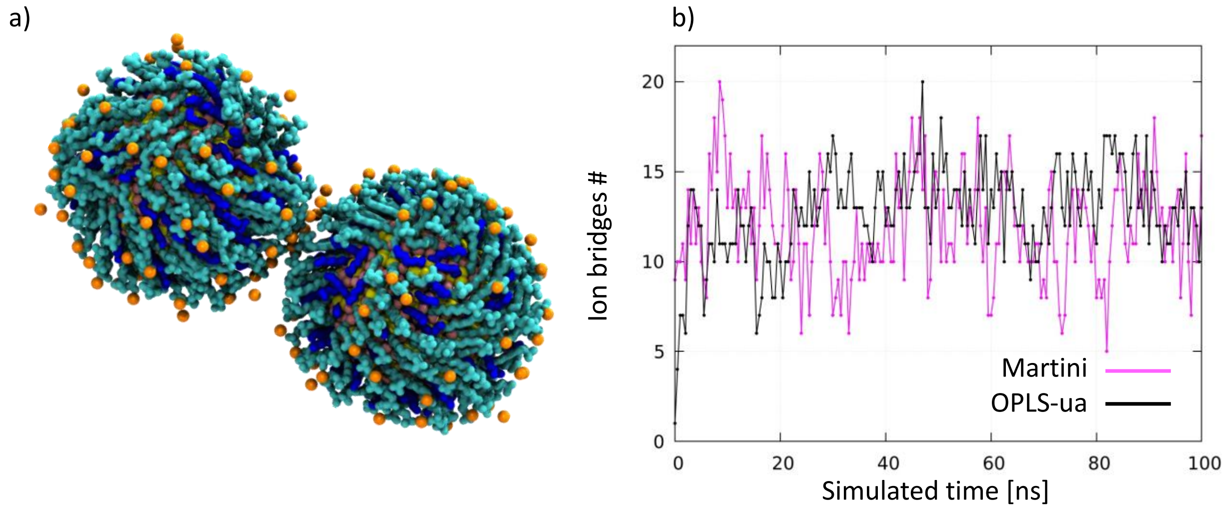


**Figure S3: OPLS dimer.** We ran 4 simulations (4x200 ns) of a dimer solvated in water with the OPLS-ua force field. We tuned the salt concentration (150 mM and 300 mM) and the charge of the counterions ($Na^+$ or $Ca^{2+}$). The dimer was stabilized by combining 300 mM NaCl and $Ca^{2+}$ counterions. a) Snapshot of the dimer in the OPLS NaCl 300 mM and $Ca^{2+}$ counterions simulation. The NP core is represented in yellow (Au) and pink (S), the ligands cyan (MUS) and blue (OT) and the $^{Ca2+}$ counterions in orange. At the interface, it is possible to see several ions bridging the MUS ligands of different NPs. b) Comparison between the number of ion bridges present at the NP-NP interface, in Martini and the OPLS-ua unbiased simulation.

**Martini limitations and comparisons with OPLS-ua.** It is known[18,19] that the use of the Martini force-field implies significant intrinsic approximations to the behavior of ions and charged moieties in general. Electrostatic interactions are treated with a cut-off to Coulomb interactions to 1.2 nm. This is expected, in general, to cause a poor description of long range screening effects. In our case, the cut-off to electrostatic interactions can reduce NP-NP electrostatic repulsion, causing an underestimation of the barrier that separates the ion-bridged dimer from the hydrophobic-contact dimer. The mechanism of ion-bridging, per se, should not be affected by the electrostatic cut-off, being a very short range effect. Another notable approximation, which Martini shares with other coarse grained models, consists in the large size of the ions which end up representing a central charge and its first hydration layer[2,20]; as a consequence of this large interaction surface, ions often behave as if they were stickier than expected. This could be in line with the observations done at united-atom resolution, where divalent ions seem necessary to induce ion-bridged aggregation. Nevertheless, the accurate description of ions is a critical aspect of both coarse grained and atomistic force fields[21,22] and here the reliability of the model is assessed mainly by comparison with the experimental data.

**Cryo-EM imaging of aggregation of MUS:OT NPs on DOPC liposomes.** The data reported in Figure S4 are taken from ref.[23] We recall here the most important details of the experimental procedures.

*NP synthesis.* Gold nanoparticles were synthesized according to the procedure described in our previous publications[9,10]. The size distribution of the nanoparticles was determined by transmission electron microscopy (TEM) and the ratio between MUS and OT was determined with nuclear magnetic resonance spectroscopy ($^1$H-NMR), as described in[9,10].

*Liposome preparation and incubation with NPs.* Large DOPC liposomes (> 100 nm) were obtained via extrusion. For cryo imaging ˜2.5 mM of vesicles were incubated with a final concentration of 0.28 mg/ml or 1.4 mg/ml nanoparticles at a final volume of 30 µl at room temperature overnight, without any agitation.

*Cryo-EM.* Grids for cryo-EM and tomograms were prepared in a commercial vitrification system (Vitrobot Mark IV, FEI, Netherlands) with 100% humidity at 22 ˚C. 4 µl of sample was pipetted on the lacey carbon grid (300 mesh, Electron Microscopy Science, Hatfield, PA) which was glow discharged for 3 seconds beforehand. Prior to plunge freezing excess sample was blotted under a blotting force of -15 for 2 seconds. After plunge freezing the grids were transferred at -178 ˚C into a Gatan 626 cryo-holder (Gatan inc. Warrendale, PA) and imaged in a FEI Tecnai F20 microscope (FEI) operated at 200kV. The total electron dose for one image was ˜30 e/Å2 and it was acquired using magnification of 50000X or 62000X (pixel size 0.2 nm and 0.16 nm respectively) with defocus values of -0.8 to -2.4 µm. Tomograms were aligned and reconstructed from a tilt series which were mostly acquired between ±60˚ with 2 degree increments. Total electron dose for the tomogram ranged from 30 to 100 e/Å2 and they were acquired using magnification 2900X (0.32 nm) and 50000X (pixel size 0.2 nm) with the defocus value ranging between -3 to -5 µm. Images for tomograms were recorded using a FalconIII camera (4096 x 4096 pixels, FEI), unless otherwise specified. Alignments of the images and reconstructions of the tomograms were done using 3DInspect (FEI).

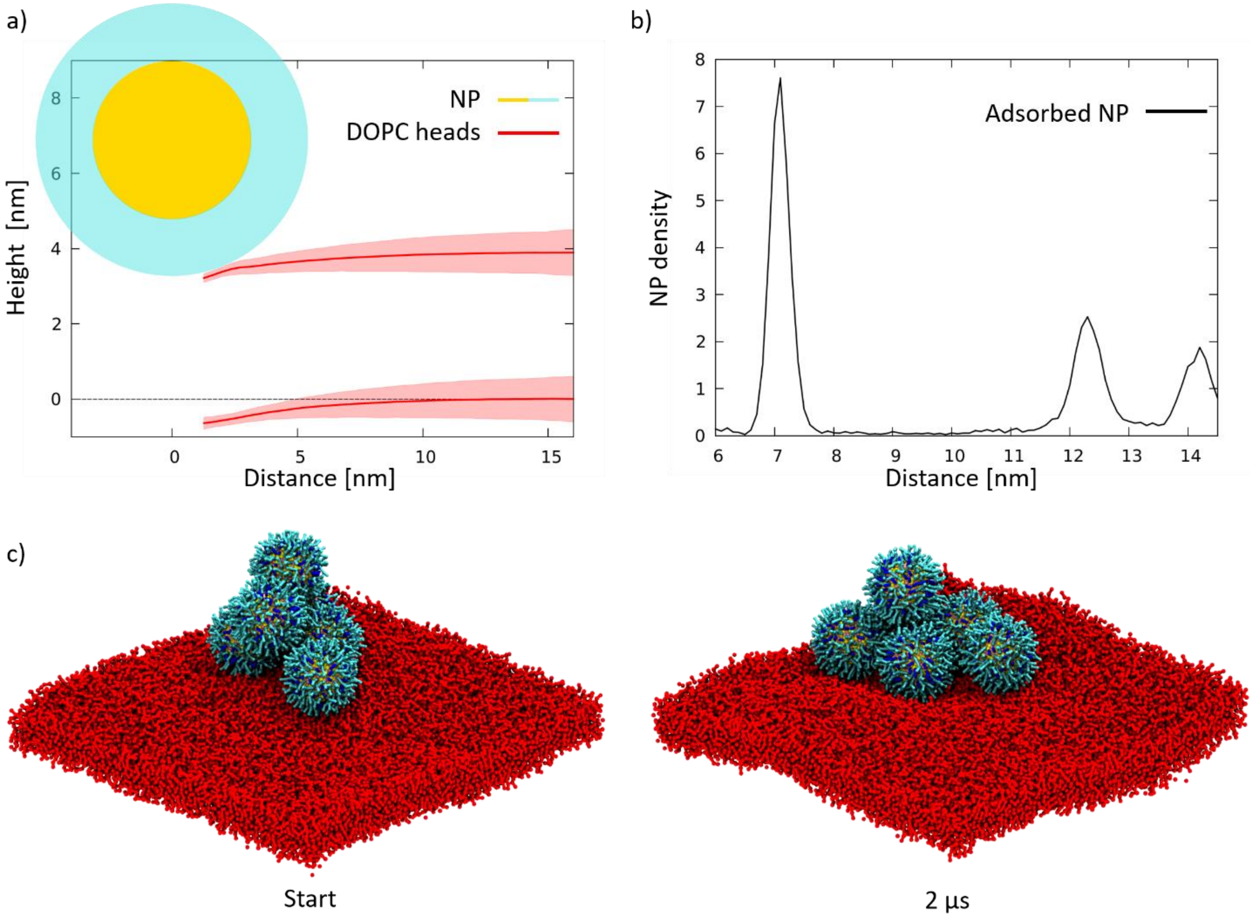


**Figure S4: Adsorbed NPs.** a) Radial profile of the DOPC lipid heads (red) in the proximity of the NP-membrane interaction site. The NP is represented with a sketch, the yellow circle being the core and the cyan circle the ligand shell. While the membrane bends in the presence of the NP, the deformation is under 15% of the thickness of the bilayer, hence there is no wrapping of the NP. b) NP-NP RDF from the 9 adsorbed NP unbiased simulation. The peak of NP-NP distance is the same of that from the simulations in water, as shown in Figure S1a. c) Flattening process: in the initial configuration, a cluster of 6 NP is adsorbed on the membrane, forming two layers of NPs: 4 NPs in the bottom layer, in direct contact with the membrane, 2 NPs in the top layer. After 2 µs one of the NPs from the upper layer manages to make contact with the membrane, and in this position it remains stable for the rest of the simulation (10 µs).

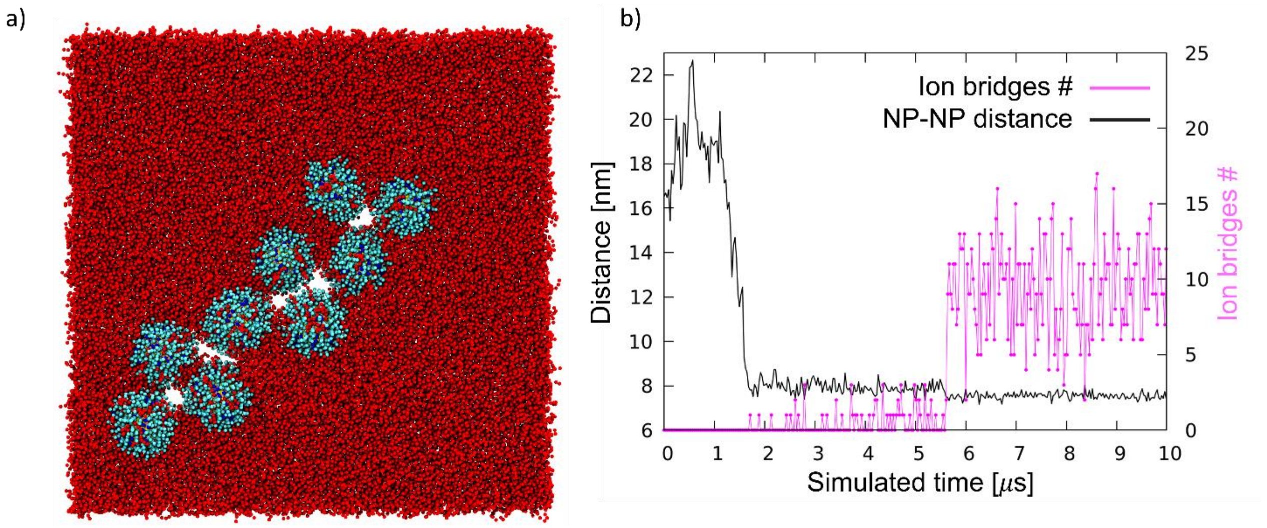


**Figure S5: Snorkeled NPs.** a) Snapshot from the unbiased simulation of the diffusion of 9 fully-snorkeled NPs. The simulation show pores formation, which are quite dynamic in shape and relative position, but correlate well to the formation of ion bridges.

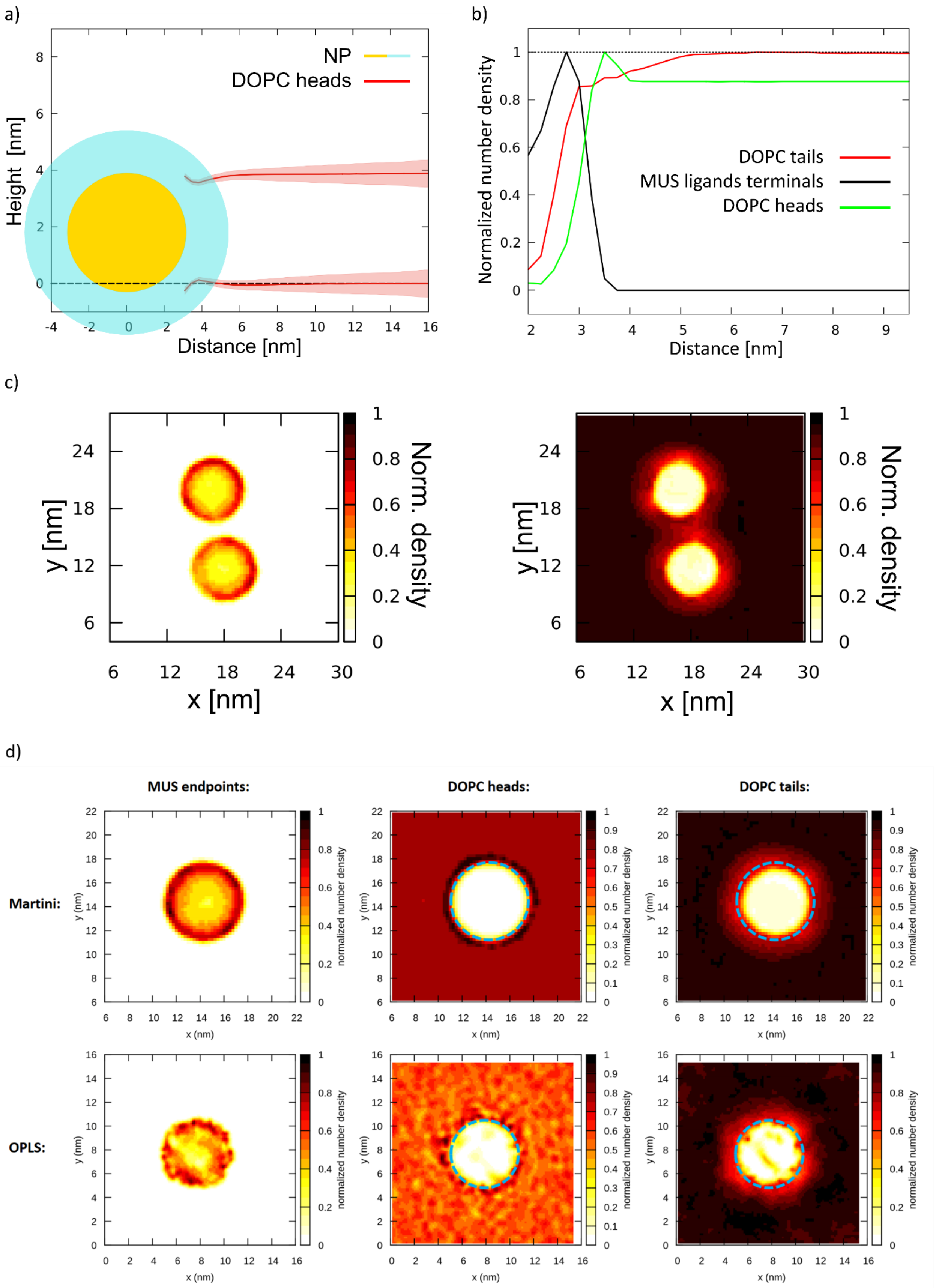


**Figure S6:** a) fully-snorkeled NPs do not alter membrane curvature. b) The lipid depletion aura is also observed in the neutralized system (n-MUS:OT NPs). c) Normalized number density maps of MUS ligands (left) and DOPC tails (right) around a fully snorkeled n-MUS:OT NPs dimer. d) Normalized number density maps for single MUS-OT NPs fully snorkeled in DOPC, showing the agreement between the CG and ua approaches.

(17) Lavagna, E.; Zeyike Pelin Güven; Bochicchio, D.; Olgiati, F.; Stellacci, F.; Rossi, G. Amphiphilic Nanoparticles Generate Membrane Curvature and Shape Liposome-Liposome Interfaces. 2021.

(23) Zekiye Pelin Güven. PhD thesis, 2018, Nanoparticle Liposome Interactions.
